# ARCS: Assembly Roundup by Chromium Scaffolding

**DOI:** 10.1101/100750

**Authors:** Sarah Yeo, Lauren Coombe, Justin Chu, René L Warren, Inanç Birol

**Affiliations:** BC Cancer Agency, Genome Sciences Centre, Vancouver, BC V5Z 4S6, Canada

**Author notes:** Authors contributed equally (Sarah Yeo, Lauren Coombe, René L Warren).

**Keywords:** 10X Genomics Chromium, ARCS, ABySS, next-generation sequencing, *de novo* assembly, genome scaffolding, read clouds

## Abstract

Sequencing of human genomes is now routine, and assembly of shotgun reads is increasingly feasible. However, assemblies often fail to inform about chromosome-scale structure due to lack of linkage information over long stretches of DNA – a shortcoming that is being addressed by new sequencing protocols, such as linked reads from 10X Genomics. Here we present ARCS, an application that utilizes the barcoding information contained in linked reads to further organize draft genomes into highly contiguous assemblies. We show how the contiguity of an ABySS *H. sapiens* genome assembly can be increased over six-fold using moderate coverage (25-fold) Chromium data. We expect ARCS to have broad utility in harnessing the barcoding information contained in Chromium data for connecting high-quality sequences in genome assembly drafts. Availability: http://www.bcgsc.ca/platform/bioinfo/software/arcs

**Supplementary information** available online.

## 1 Introduction

The Chromium sequencing library preparation protocol from 10X Genomics (10XG, Pleasanton, CA) builds on the Illumina sequencing technology (San Diego, CA) to provide indexing/barcoding information along with short reads to localize the latter on long DNA fragments, thus benefiting the economies of scale of a high-throughput platform. As sequence reads from 20 to 200 kb molecules are barcoded/linked, applications of the technology has mainly focused on phasing variant bases in human genomes ([1] and [2]). The ability to generate linked reads with 10XG is akin to that of Illumina TruSeq [3]. The latter technology provides useful complementary information to whole genome shotgun assembly projects, as the pseudo-long reads it generates may help resolve long repeats. However, to generate pseudo-long reads, TruSeq requires high coverage data of the co-localized reads for a priori fragment assemblies (by default, transparent to the user), essentially generating low fragment coverage data for its target genome. Hence, TruSeq may be prohibitively expensive for providing mammalian-sized genomes with adequate fragment coverage. Conversely, the Chromium platform typically provides low-coverage for each single barcoded molecule, limiting its utility for individual fragment assembly. Though, it makes up for this limitation in throughput, providing higher fragment coverage.

Recently this data type has been utilized for scaffolding a draft genome assembly [4], using a software designed to scaffold sequences using contiguity preserving transposition sequencing (CTP-seq) and Hi-C data, another long-range information data source [5]. In their paper, Mostovoy and colleagues [4] showed 12-fold improvement in contiguity of a human genome assembly draft using GemCode sequencing (precursor to Chromium, from 10XG) at 97-fold coverage, demonstrating the potential of the technology for scaffolding draft genomes. Here we present ARCS, the Assembly Round-up by Chromium Scaffolding algorithm, a method that leverages the rich information content of high-volume long sequencing fragments to further organize draft genome sequences into contiguous assemblies that characterize large chromosome segments. We use the recent Genome In A Bottle (GIAB) human genome sequence data [6], and compare ARCS to fragScaff, the only other technology shown in a publication to utilize 10XG linked reads for scaffolding genome assembly drafts [4]. We also present similar benchmarks to Architect, a recently published scaffolder [7] shown to work on Illumina TruSeq synthetic long sequences’ underlying short reads (read clouds) and suggested to be adaptable to Chromium data. We show how our implementation yields assemblies that are more contiguous and accurate than fragScaff and Architect over a wide range of parameters while using less time and compute resources. We note that ARCS scaffolding of pre-existing human genome drafts using two different linked-reads datasets yields assemblies whose contiguity and correctness is on par with or better than those assembled with newly released 10XG Supernova *de novo* assembler [8].

## 2 Methods

We used two human Chromium read datasets for scaffolding, including one from individual NA12878 and GIAB HG004 NA24143. For the genome assembly contig and scaffold baselines, we first downloaded Illumina whole genome shotgun (WGS) 2x250 bp paired-end sequencing data and Illumina 6 kbp mate-pair data for an Ashkenazi female (NA24143). ABySS-2.0 [9] was used to assemble the whole-genome shotgun paired-end and mate-pair reads into contigs and scaffolds and Chromium reads were aligned to those sequences. Using these alignments as input, we ran and benchmarked ARCS (v1.0.0), Architect (v0.1) and fragScaff (v140324) to further scaffold contig and scaffold baseline sequences 3 kbp and longer, as recommended [4], while investigating the effects of multiple parameter combinations on scaffolding (Table S1), reporting contiguity length metrics and breakpoints from sequence alignments to the reference human genome, which serves as a proxy for counting large-scale misassemblies in both contigs and scaffolds.

### 2.1 ARCS algorithm

The pipeline collectively referred to as ARCS first pairs sequences within a draft assembly, then lays out the pairing information for scaffolding. In the sequence pairing stage (Fig. 1), input alignments in BAM format are processed for sets of read pairs from the same barcode that align to different sequences. These form a link between the two sequences, provided that there is sufficient number of read pairs aligned (-*c*, set to 5 by default). Each link represents evidence that one barcode/molecule connects both sequences. To account for barcode sequencing errors, only barcodes within a specified multiplicity range (-*m*) are considered (default 50-10000 for Chromium). The multiplicity refers to the read frequency of each barcode and the range, a specific slice of reads that are considered by ARCS. In our experience, this distribution is wider for GemCode and should be set to 50-10000. As we are interested in ordering and orienting sequences, we consider reads that align to the 5’ and 3’ end (-*e*) bases of each sequence. This parameter effectively sets the maximum length of regions, at the end of sequences, where Chromium reads align. Reads whose alignments fall outside of these regions are not considered and thus, adjusting -*e* to a lower value to account for shorter contigs is important as it will have for effect to mitigate ambiguity when creating an edge between any two sequences. In addition, as the BAM file is read, only reads that align to a sequence with at least the specified sequence identity (-*s*, set to 98 by default), map in proper pairs and are unique alignments are considered. This ensures that only high-quality alignments provide evidence for the subsequent linking stages, as alignments involving reads with long repeat regions or chimeric reads will be skipped.

**Fig. 1.**
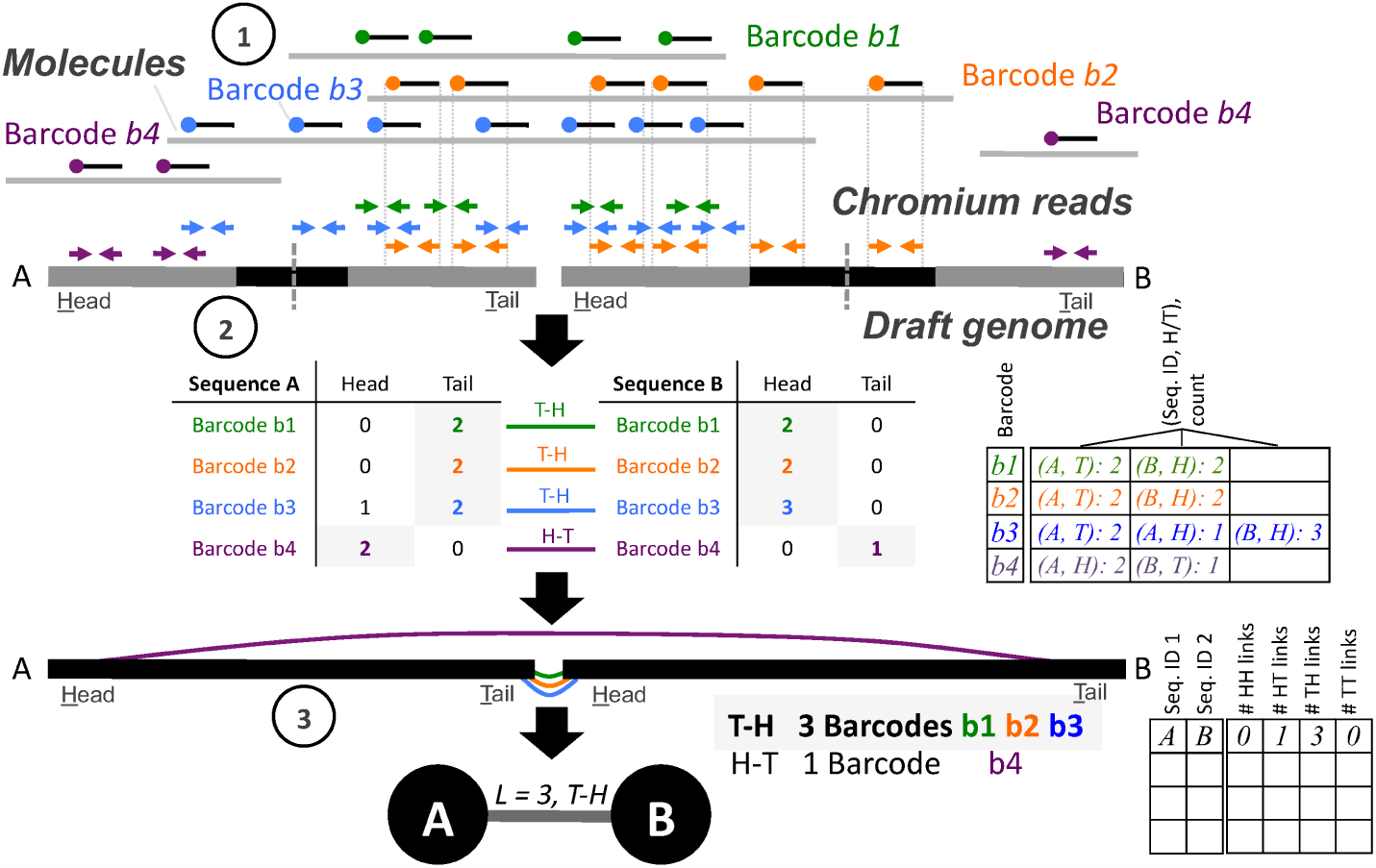
ARCS algorithm. 1) 10XG Chromium reads (blue, green, orange, and purple arrows) are aligned to the draft genome. 2) Sequences are split in half by length and the ends of each are considered the head (H) or tail (T) regions (represented with grey boxes, length of the ends controlled by ARCS parameter -e). The number of read pairs derived from the same barcode and aligning to the head (H) or tail (T) regions of the sequence are tallied. These tallies are stored in memory using a map data structure, where the key is the barcode sequence. The value maps a tuple of the sequence ID and H or T to the count of the number of read pairs of that barcode which map to the corresponding region of the sequence. 3) The number of barcodes supporting each link orientation (H-H, H-T, T-H, T-T) between sequence pairs is tallied. The tallies are stored in memory using a map data structure, where the key is a pair representing the two potentially linked sequences, and the value is a vector of integers representing the number of barcodes supporting each possible link orientation. For a given barcode to contribute linking evidence, the distribution of reads of that barcode aligning to the H or T regions of both sequences in the potential pair must significantly differ from a uniform distribution. A dot file is then generated which encodes the linkage evidence, where links (edges) between two sequences (nodes) are only added if the link orientation with the maximum support is predominant.

The relative orientations of sequences are inferred through the read alignment positions. First we determine, using read alignments, subsets of reads with the same barcode that co-locate within one end of the sequence (Fig. 1, step 1); Each sequence is split in half by length and one end region arbitrarily labeled head (H), the other tail (T). The number of read pairs of the same barcode aligning to the head or the tail of a sequence (within -*e* bp or less of the end) is tallied as the BAM file is read. A map data structure tracks, for a given barcode, the number of reads that map to the H or T of a sequence (Fig. 1, step 2). Once the alignment file is read into memory, every possible pair of sequences which have a sufficient number of aligned reads from a given barcode (-*c*) are considered. For each sequence in a potential pair, a binomial test is used to calculate whether the observed distribution of reads aligning to the 5 or 3 end of a sequence is significantly different from a uniform distribution (threshold *p*=0.05, parameter -*r*). Likewise, the number of linking barcodes/indices that support each of the four link orientations (H-H, H-T, T-H, T-T) is tallied in a map data structure for each potential sequence pair (Fig. 1, step 3). Using the link orientation tallies for each sequence pair, a graph data structure is constructed, where the nodes are sequences, and the edges represent links between them. An edge is formed only if the link orientation, defined by the order of sequence pairs head and tail regions, is the most represented combination across supporting barcodes.

After the sequence pairing stage completes, ARCS outputs a single file in the graph description language (gv) format. The gv file is converted using supplied python scripts (makeTSVfile.py), to a tab-separated value (tsv) file listing all possible oriented sequence pairs, the number of supporting barcodes with gap sizes arbitrarily set at 10 bp. Since positional information of reads within the molecule of origin is not known, estimates of gap sizes are not possible. At the layout stage, the latter tsv file is read and scaffolds constructed using the algorithm implemented in LINKS (v1.7 and later), as described [10]. Because linked sequence pairs may be ambiguous (a given sequence may link to multiple sequences), sequences are joined only if the number of links connecting a sequence pair is equal or greater than a minimum (LINKS parameter -*l*, default of 5). Ambiguous pairings are resolved when the ratio of barcode links of the second-most to top-most supported edge is equal or below a threshold (LINKS parameter -*a*, default of 0.3; we recommend higher values such as -*a* 0.7 and 0.9 when running LINKS within ARCS).

ARCS is implemented in C++ and runs on Unix.

### 2.2 Data sources

Two human Chromium datasets were downloaded, including one from individual NA12878 and GIAB HG004 NA24143 (Table S2). The Illumina WGS 2x250 bp paired-end sequencing data of an Ashkenazi mother (NA24143) was downloaded from https://github.com/genome-in-a-bottle/giab_data_indexes/blob/master/AshkenazimTrio/sequence.index.AJtrio_Illumina_2x250bps_06012016 under accession number NIST HG004 NA24143 SRS823307.

The Illumina 6 kbp mate-pair sequencing data from that individual was downloaded from https://github.com/genome-in-a-bottle/giab_data_indexes/blob/master/AshkenazimTrio/sequence.index.AJtrio_Illumina_6kb_matepair_wgs_08032015.

We downloaded the corresponding 10X Genomics 2x128 bp barcoded paired-end sequencing data for that individual [6] from https://github.com/genome-in-a-bottle/giab_data_indexes/blob/master/AshkenazimTrio/alignment.index.AJtrio_10Xgenomics_ChromiumGenome_GRCh37_GRCh38_06202016.

The 156-fold coverage NA12878 10XG Chromium data was downloaded from: http://support.10xgenomics.com/de-novo-assembly/datasets/msNA12878, which we assembled with the 10XG Supernova v1.1 *de novo* assembler (available at http://support.10xgenomics.com/de-novo-assembly/software/downloads/latest), as described [8].

### 2.3 Data analysis

Prior to assembly, the adapters from the mate-pair reads were removed using NxTrim v0.4.0 [11] (with parameters -*norc -joinreads -preserve-mp*), which also classifies reads as mate-pair, paired-end, single-end or unknown. Only reads classified as mate-pair were used for scaffolding of contigs with ABySS. Both paired-end and mate-pair reads were corrected with BFC v181 [12] (with the parameter-s3G). Contigs and scaffolds were assembled with ABySS v2.0 [9] with the command: abyss-pe *name*=hsapiens *np*=64 *k*= 144 *q*=15 *v*=-v *l*=40 *s*=1000 *n*= 10 *S*=1000-10000 *N*=7 *mp6k_de*=–mean *mp6k_n*= 1 *lib*=pe400 *mp*=mp6k, where pe400 and mp6k are variables listing all files containing paired-end sequencing and MPET reads, respectively.

The 10XG Chromium sequencing data was converted from a BAM file to a FASTQ format. Briefly, the read barcodes were extracted from the RX tag in the BAM file and appended to the read name following an underscore character (eg. NAME_BARCODE). Read pairs were only added to the FASTQ file if the corresponding alignment had a sequence identity of at least 90% and were aligned in proper pairs. Chromium reads were then aligned to the contigs and scaffolds using BWA mem v0.7.13 (default values, adjusting *t* for multiple threads) [13] and sorted by name.

Based on recommendations, the input to fragScaff also included a n-base bed file, generated by the provided script fasta_make_Nbase_bed.pl, and a repeat bed file generated by performing a blastn v2.4.0 alignment [14] of the input assembly to itself (with parameters *word_size* 36, *-perc_dentity* 95, *-outfmt* 6), which was then processed by the provided script blast_self_alignment_filter.pl. The scripts that ran on the data described above are available at ftp://ftp.bcgsc.ca/supplementary/ARCS, along with the commands and parameters we ran and set with each tool and corresponding assemblies.

The NG50 and NGA50 length metrics reported were calculated using a genome size of 3,088,269,832 bp.

Benchmarking was done on a DELL server with 128 Intel(R) Xeon(R) CPU E7-8867 v3, 2.50GHz with 2.6TB RAM.

## 3 Results

### 3.1 Scaffolding with the NA24143 GIAB Chromium data

We measured the contiguity (NG50 and NGA50 length metric) and correctness of resulting assemblies after ARCS, Architect and fragScaff scaffolding of baseline assemblies (Table S3). During this process, we tested the effect of various parameters, including the scaffolding-specific -*a*, -*u* and –*rc-rel-edge-thr* (abbreviated rel) parameters in the corresponding tools. Generally, these parameters affect scaffolding stringency by evaluating the validity of the linkages. Mostovoy *et al*. [4] reported their best assembly using fragScaff parameters -*j*1 and -*u*3, prompting us to explore similar values of -*j* and -*u* on our dataset. These parameters are described as *the mean number of passing hits per node to call the p-value cutoff* and *modifier to the score to consider the link reciprocated* in fragScaff and are the parameters previously optimized [5]. In Architect, the parameters -*rc-abs-thr*, -*rc-rel-edge-thr* and -*rc-rel-prun-thr* control the minimum number of reads from a given barcode aligning to sequences required to create an edge in the scaffold graph, the relative barcode support needed for creating edges and the relative barcode support needed for pruning edges, respectively.

To assess correctness, we aligned the assemblies to the primary chromosome sequences of the human reference GRCh38 and counted the number of observed breakpoints using abyss-samtobreak [9]. At the contig level (Fig. 2a), we observe that while the ARCS and fragScaff assemblies (highest contiguity achieved at -*a*0.9 and -*u*2, in that order) have similar sequence contiguity (NG50 of 303,034 vs. 304,926 bp, respectively), the ARCS assembly has less than one third the number of breakpoints compared to fragScaff (2,030 vs. 6,345). In context, the corresponding ARCS and fragScaff assemblies have 16.3% and 263.4% more breakpoints than the baseline contig assembly, respectively (Tables S3-S5). This indicated that, while the resulting fragScaff assemblies were highly contiguous, they might harbor substantially more misassemblies. Architect scaffolding of the baseline contig assembly did not yield appreciable gains despite extensive parameter tuning (Fig. 2a and Table S6). Contig-level scaffolding took more than 5 hours and over 5 days for fragScaff and Architect, respectively (Table 1; complete benchmark results are available at: ftp://ftp.bcgsc.ca/supplementary/ARCS/benchmarks).

**Fig.2.**
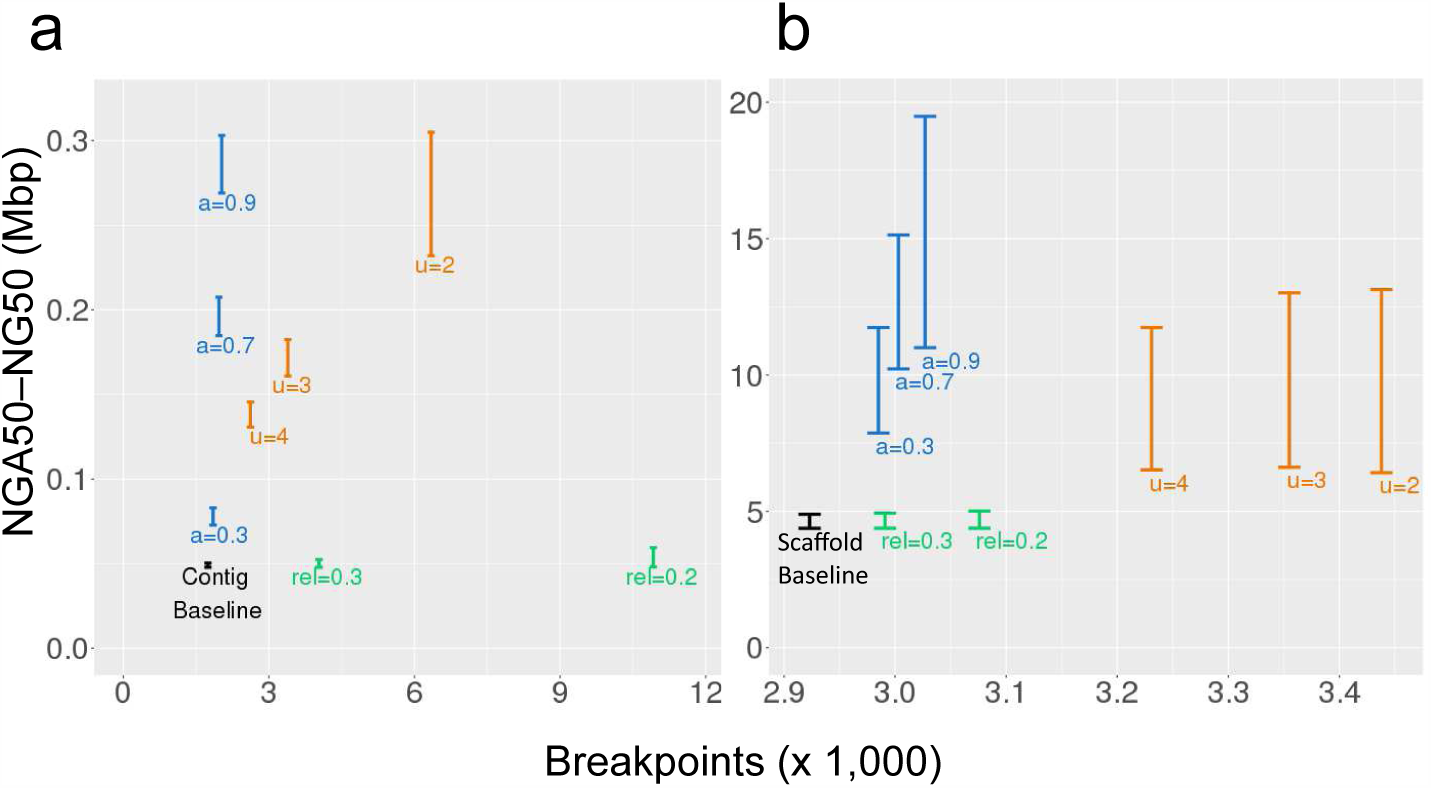
Contiguity and correctness resulting from scaffolding (a) contig or (b) scaffold baseline assemblies with 10XG Chromium reads using fragScaff (orange), Architect (green), and ARCS (blue). We show the effect of the scaffolding parameters -*u* (fragScaff), –*rc-rel-edge-thr* (abbreviated rel, Architect) and -*a* (ARCS). The Y-axes show the range of NGA50 to NG50 lengths to indicate the uncertainty caused by real genomic variations (captured by breakpoints analysis) between individual NA24143 and the reference genome GRCh38. The X-axes show the number of breakpoints that occur when aligning the resulting assembly to the reference.

**Table 1.** Total wall-clock time and peak memory usage for ARCS (-*c*5 -*e*30000 -*r*0.05 -*l*5 -*a*0.3), Architect (-*t*10 or -*t*5 -*rc-abs-thr*5 –*rc-rel-edge-thr*0.4 –*rc-rel-prun-thr*0.2) and fragScaff (-*C*5 -*E*30000 -*j*1 -*u*4) scaffolding applied to baseline contig and scaffold assemblies.

| Scaffolder | Baseline assembly | Number of threads | Wall-clock time (h:mm) | Peak memory (GB) |
| --- | --- | --- | --- | --- |
| ARCS | contig | 1 | 1:12 | 9.4 |
| Architect | contig | 1 | 141:18 | 11.1 |
| fragScaff | contig | 64 | 6:37 | 8.1 |
| ARCS | scaffold | 1 | 0:55 | 3.4 |
| Architect | scaffold | 1 | 6:07 | 9.6 |
| fragScaff | scaffold | 64 | 1:59 | 14.1 |

At the scaffold level (Fig. 2b), we observe that ARCS achieves a greater sequence contiguity and correctness than Architect and fragScaff (NG50 (Mbp) / breakpoints, 19.5 / 3,027 vs. 5.0 / 3,076 vs. 13.1 / 3,438 in that order) when comparing amongst the most contiguous assemblies for each tool (Tables S4-S6). This is despite there being roughly one order of magnitude fewer misassemblies between fragScaff and ARCS assemblies that used the scaffold as opposed to contig baseline sequences for scaffolding (411 vs. 4,315 breakpoints, respectively). These assemblies respectively harbor 3.6%, 5.2% and 17.6% more breakpoints than the baseline scaffold assembly, which suggests that ARCS and other scaffolders for 10XG data work best when the draft to re-scaffold is more contiguous. To see whether these 411 additional breakpoints in the fragScaff vs. ARCS assemblies are large-scale misassemblies, we aligned the corresponding assemblies to the reference human genome and plotted the alignments (Fig. 3). As observed, fragScaff scaffolding of the baseline scaffold sequences yields more inter-chromosomal translocation misassemblies (Fig. 3a) when compared to ARCS (Fig. 3b). We note that increasing the fragScaff -*j* parameter (mean passing links across nodes) while relaxing -*u* (score cut-off multiplier) yields assemblies whose contiguity rival that of ARCS (16.9 vs 19.5 Mbp NG50, respectively), but at the cost of increased misassemblies (Tables S4 and S5). Architect scaffolding of both contig and scaffold baseline assemblies yielded a marginal increase in contiguity figures (Fig. 2 and Table S6), which we can only speculate on. Apart from the tool design, which is intended for Illumina TruSeq read clouds [7], it is possible that the sequencing depth of the 10X Genomics linked reads used in our study (25-fold) may not be adequate to observe appreciable gains with this tool.

**Fig. 3.**
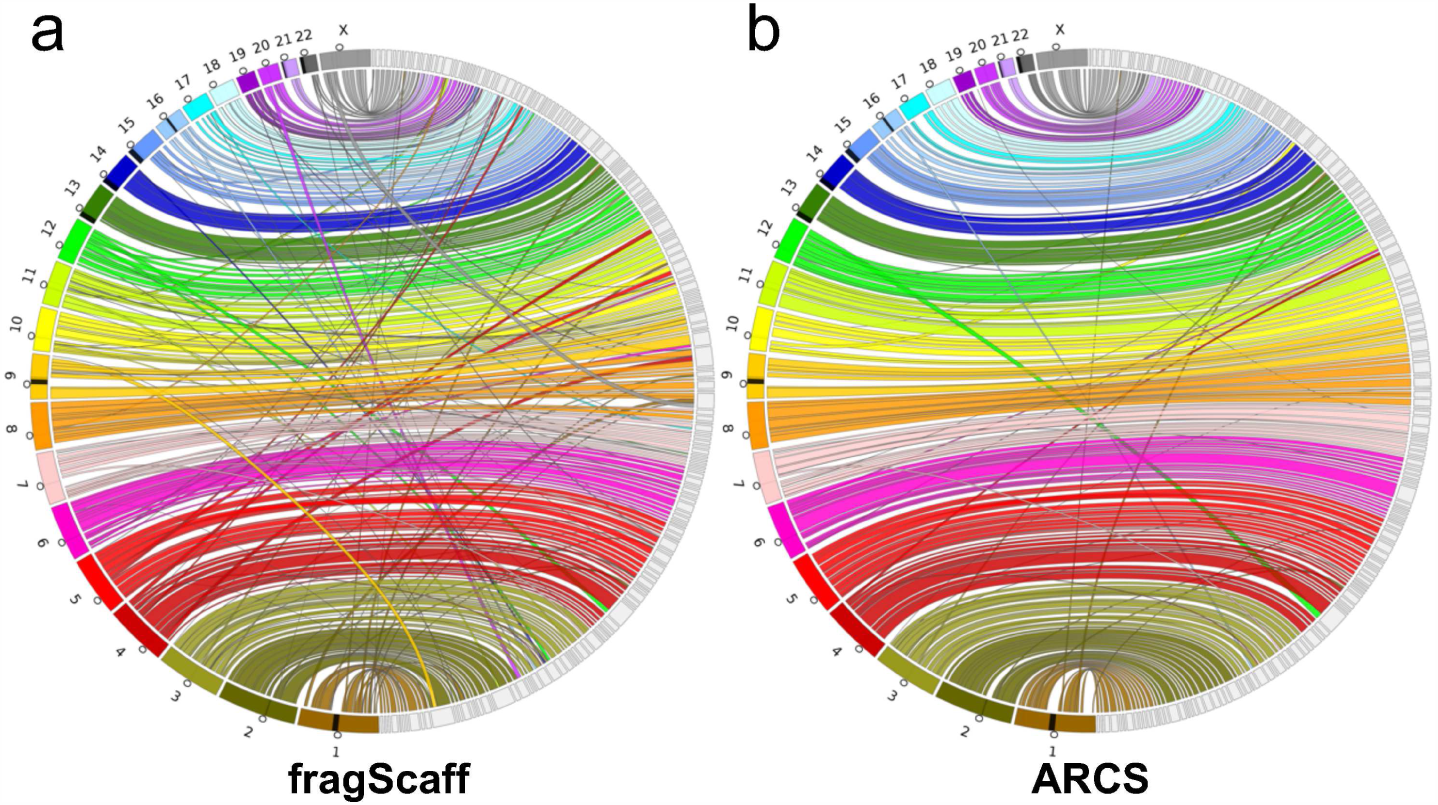
(a) A Circos [15] assembly consistency plot of conservative fragScaff (-*C*5 - *E*30000 -*j* 1 -*u*4) and (b) ARCS (-*c*5 -*e*30000 -*r*0.05 -*l*5 -*a*0.3) scaffolding of the baseline scaffold assembly. Scaftigs from the largest 175 (fragScaff) and 177 (ARCS) scaffolds, consisting of 75% of the genome are aligned to GRCh38 with BWA mem. GRCh38 chromosomes are displayed on the left while scaffolds are displayed on the right. Connections show the aligned regions between the genome and scaffolds. The open circles along chromosomes indicate the centromeres, while the black regions on chromosomes indicate gaps in the reference.

We also compared the resource efficiency of all three tools over the parameter range tested (See ftp://ftp.bcgsc.ca/supplementary/ARCS/benchmarks) and report its runtime and memory usage on the most contiguous assemblies of baseline scaffold sequences (Table 1). ARCS outperforms Architect and fragScaff for run time on both contigs and scaffolds (average 3 fold faster than fragScaff) and memory usage on scaffolds (4 times less memory when compared to fragScaff). It should be noted that the run time of Architect and fragScaff increases quadratically with the number of input sequences, making them inefficient choices for assemblies with a large number of input sequences (more than 250,000). Running Architect on the baseline contig assemblies took roughly 6 days (141 h) for most parameter combinations. In contrast, equivalent runs of this tool on the scaffold baseline assemblies were faster (6 h) due to having 20 times less sequences to process (Table S3). The execution speed of ARCS on the contig and scaffold baseline assemblies was consistent, both finishing in approximately 1 hour (1 h 12 m and 55 m, respectively).

### 3.2 Scaffolding with the NA12878 Chromium data

Recently, 10XG released their *de novo* assembly software called Supernova, which implements a scaffolding stage and is developed specifically for assembling Chromium data [8]. The authors presented a variety of human genome assemblies, each yielding N50 contiguity lengths 15 Mbp or higher, factoring in scaffolds 10 kbp and larger. We re-capitulated the Supernova experiment on 156-fold Chromium sequencing data for the NA12878 individual (Table S2), and corroborate their results (Table S7). When applying a scaffold sequence length cut-off on par with that used in our study (500 bp), we report N50 length metrics corrected for genome size (NG50) in the megabase range (14.7e6 bp), which is consistent with what was maximally achievable with ARCS using either the same dataset (NG50=18.3e6 bp) or the lower-coverage GIAB Chromium dataset (NG50=19.5e6 bp) that used the same scaffolding parameters. A Supernova assembly of 51-fold raw GIAB chromium reads produced a similarly contiguous assembly (NG50=13.5e6 bp).

Despite the NA12878 Chromium read data used with ARCS having substantially deeper coverage (5x deeper, Table S7 datasets 5 vs. 3), we observe that ARCS performs consistently across both human NA12878 and NA24143 datasets. Perhaps more interesting is the observation that there are only marginal gains in N50 length contiguity when using the higher coverage Chromium dataset (21.8e6 vs. 22.2e6 bp when using NA24143 GIAB vs. NA12878 data, with parameters -*e* 30,000 -*r* 0.05 -*c* 5 -*l* 5 -*a* 0.9) in spite of making 406 additional merges (Tables S2 and S7). This indicates that, under the conditions tested herein, with the draft assembly utilized and parameters set, the solution may work optimally with less data. We do stress the importance of knowing the distribution of reads within each barcode as it may vary between datasets, which is what is observed for NA24143 and NA12878 (Fig. S1). In our experience, this distribution is wider for GemCode and should be set to 50-10000 to include the bulk of the data while discarding outliers.

While we show that the contiguity of a *H. sapiens* contig assembly can be increased over six-fold with the use of 10XG data with only a marginal increase in probable errors with an average ± S.D. of 196 ± 77 total breakpoints compared to the baseline contig assembly, there are limitations to the method. As mentioned by Adey *et al.* [5], when using a barcode-based approach, it is difficult to confidently place short input sequences due to a lower number of indexed read pools aligning to the sequence. In addition, as the barcoded molecules may be over 100 kbp in length [16], it is possible that they span several entire short input sequences, preventing ARCS from extracting orientation information from read alignment positions, as they do not preferentially align to one end. Barcode reuse across molecules or incorrect alignment of linked reads due to repeats can also introduce false linkages at the sequence pairing stage, resulting in incorrect merges during the scaffolding stage.

With Chromium, 10X Genomics improved upon the GemCode protocol by increasing the number of fragment partitions and curbing barcode reuse. While the positional information of linked reads within a given fragment is not known, making it challenging for estimating gap or overlap sizes in genome assemblies, it remains an attractive technology for scaffolding draft genomes. This is especially true when the technique is applied to later stages of scaffolding, when the contiguity of the draft sequence assembly is high. To our knowledge, ARCS is the first publicly available stand-alone application for scaffolding draft genomes that is designed specifically for 10X Genomics Chromium reads.

## Acknowledgements

This work has been supported by the National Human Genome Research Institute of the National Institutes of Health (under award number R01HG007182), with additional support provided by Genome Canada, Genome British Columbia, and the British Columbia Cancer Foundation. The content is solely the responsibility of the authors, and does not necessarily represent the official views of the National Institutes of Health or other funding organizations.

